# Sentinel cells enable genetic detection of SARS-CoV-2 Spike protein

**DOI:** 10.1101/2021.04.20.440678

**Authors:** Zara Y. Weinberg, Claire E. Hilburger, Matthew Kim, Longxing Cao, Mir Khalid, Sarah Elmes, Devan Diwanji, Evelyn Hernandez, Jocelyne Lopez, Kaitlin Schaefer, Amber M. Smith, Fengbo Zhou, QCRG Structural Biology Consortium, G. Renuka Kumar, Melanie Ott, David Baker, Hana El-Samad

## Abstract

The COVID-19 pandemic has demonstrated the need for exploring different diagnostic and therapeutic modalities to tackle future viral threats. In this vein, we propose the idea of sentinel cells, cellular biosensors capable of detecting viral antigens and responding to them with customizable responses. Using SARS-CoV-2 as a test case, we developed a live cell sensor (SARSNotch) using a de novo-designed protein binder against the SARS-CoV-2 Spike protein. SARSNotch is capable of driving custom genetically-encoded payloads in immortalized cell lines or in primary T lymphocytes in response to purified SARS-CoV-2 Spike or in the presence of Spike-expressing cells. Furthermore, SARSNotch is functional in a cellular system used in directed evolution platforms for development of better binders or therapeutics. In keeping with the rapid dissemination of scientific knowledge that has characterized the incredible scientific response to the ongoing pandemic, we extend an open invitation for others to make use of and improve SARSNotch sentinel cells in the hopes of unlocking the potential of the next generation of smart antiviral therapeutics.

## Introduction

The astoundingly rapid scientific response to the global COVID-19 pandemic is built on decades of basic scientific research. Sustained exploration of currently available therapeutic modalities as well as the generation of new treatment approaches through research investments in blue-sky technologies is necessary to combat future threats. Engineered live-cell therapies, in which cells are programmed to find sites of infection or disease and deliver precisely targeted therapeutic interventions, are promising platforms to robustly respond to a myriad of future threats.

Current cell-based therapies are primarily focused on treating cancer, although a variety of potential applications are currently under exploration^1–3^. The tools available to program these therapies are also rapidly evolving, with considerable work dedicated to improving the cellular circuits that confer targeting specificity against cancer^4–7^. Progress has been fueled by investments in synthetic biology tools that are capable of functioning orthogonally or in parallel with natural cellular systems^8–10^. A particularly exciting opportunity resides in the rise of *de novo*-designed proteins as *in silico* and on demand building blocks for cellular engineering^11–16^. Consequently, there is considerable potential in combining live cell platforms, synthetic biology and protein design to build modular and rapidly adaptable tools to combat emerging threats.

The SARS-CoV-2 virus at the root of the COVID-19 pandemic has become a proving ground for nascent technologies. At the forefront, multiple vaccines^17,18^ have been developed based on the decades-old promise of mRNA therapeutics^19^. Additionally, testing facilities have been rapidly deployed to confront growing case numbers^20^ and CRISPR-based strategies have been applied for viral detection and neutralization^21^. Even further, neutralizing proteins with potential therapeutic uses have been developed, including synthetic nanobodies^22^ and *de novo*-designed protein binders^23^. This work is a testament to decades of basic research in synthetic biology, and pioneering work in coronavirus biology that made the SARS-CoV-2 S antigen, or Spike protein, amenable to structural studies^24–27^.

Using SARS-CoV-2 as a test case, we propose a class of engineered cells capable of serving as sentinels that can detect and respond to virus-infected cells. To achieve this proof of principle, we combined the malleable synthetic Notch receptor (Syn-Notch) platform for generating customizable genetic responses to detected antigens^10,28^ with *de novo*-designed anti-Spike binders^23^ to create engineered cells that detect the presence of the Spike protein. We demonstrate that these sentinel cells can detect purified Spike protein and a model of infected cells that are expressing Spike, and we show that this sensing approach is portable between different cell types, including primary human T lymphocytes and adherent cells. We suggest that this system, when coupled with appropriate therapeutic payloads, represents a potential next-generation targeted therapeutic against future pathogenic threats.

## Results

### SARSNotch combines *de novo* designed protein binders and SynNotch to detect SARS-CoV-2 Spike

We sought to develop a sentinel cell capable of sensing the SARS-CoV-2 Spike protein, either on its own or when displayed by infected cells. Our ideal system would feature a swappable antigen-sensing domain, for use against a potentially rapidly evolving target, and a customizable genetic response. The modular SynNotch receptor^10,28^ provided a suitable platform for these requirements. A SynNotch receptor comprises an extracellular sensor domain that recognizes a desired epitope, fused to a user-defined cleavable intracellular domain that retains a transcription factor (TF) at the membrane. Antigen recognition induces the cleavage of a TF, which translocates into the nucleus to activate output expression (**Figure 1A**). Notably, Syn-Notch has previously been used to successfully detect the hepatitis B virus (HBV)^29^, thus we were optimistic that it could be adapted for use as an extracellular sensor to detect the SARS-CoV-2 Spike protein and generate a customizable transcriptional output upon this detection.

**Figure 1:**
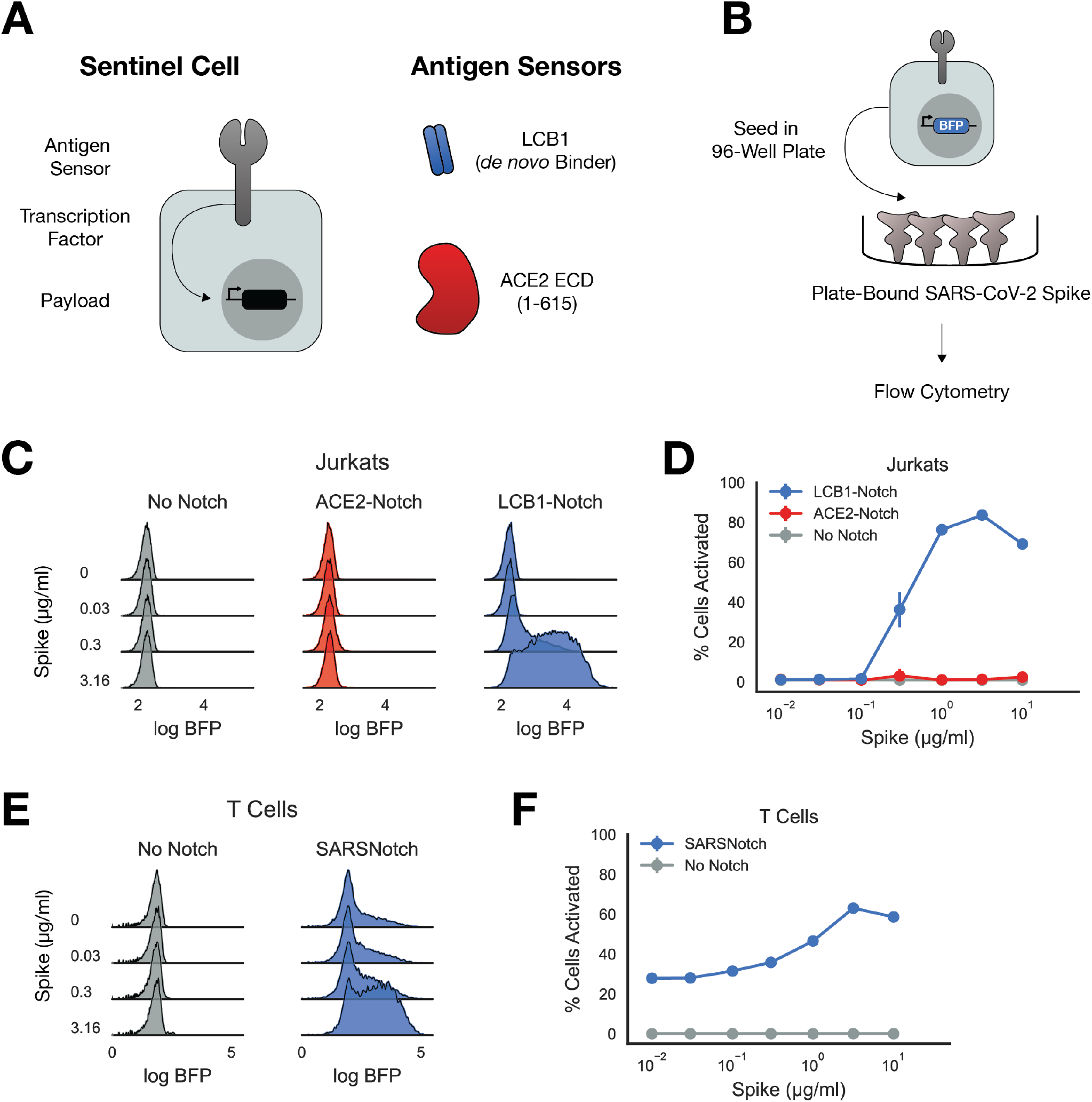
SARSNotch combines *de novo* designed protein binders and SynNotch to detect SARS-CoV-2 Spike. A) Schematic of a sentinel cell, with SynNotch’s customizable sensing, transcription, and output components, and the primary antigen sensors used in this study. B) Schematic of Spike sensing experiment. Sentinel cells expressing No Notch, ACE2-Notch, or LCB1-Notch and a BFP fluorophore downstream of SynNotch activation are incubated with purified SARS-CoV-2 Spike before being assessed for activation via flow cytometry. C) Density estimates of Jurkat sentinel activation showing logBFP fluorescence as a function of μg/ml Spike protein. D) Percent of activated Jurkat cells sentinels (assessed as described in **Supplementary Figure 1C)** plotted as a function of Spike dose. E) Density estimates of T Cell sentinels as a function of μg/ml Spike protein. F) Percent of activated T Cell sentinels plotted as a function of Spike dose (assessed as described in **Supplementary Figure 1C)**. Data points and error bars represent mean and standard deviation of 3 biological replicates.

For the transcriptional output, we used the well established Gal4p-VP64 hybrid transcription factor upstream of a promoter containing a 5X tandem GAL4 DNA activating sequence and a ybTATA core promoter ((5X)UAS-ybTATA)^28,30^. For the extracellular sensor, we initially identified Spike’s endogenous target, the extracellular domain of human ACE2^25,26^. We also explored the use of *de novo* designed binders against the Spike protein as potential antigen sensors. A computational campaign was conducted to identify short protein sequences that were able to bind the Spike protein^23^, and we incorporated the lead candidate from this screen (LCB1) into our initial antigen sensor panel (**Figure 1A**). Using our Mammalian Cloning Toolkit^31^, we rapidly constructed multiple SynNotch variants with these different extracellular sensors for Spike.

We started with a polyclonal Jurkat cell line expressing an output circuit consisting of mCitrine as a transduction marker and TagBFP downstream of (5X)UAS-ybTATA, and then transduced these cells with lentivirus encoding for either ACE2-Notch or LCB1-Notch (**Supplementary Figure 1A**). We sorted for a polyclonal cell population expressing these SynNotch variants on their surface (**Supplementary Figure 1B**). We then cultured these cells in a 96-well plate pre-coated with different concentrations of purified Spike trimer, and assayed for activation via flow cytometry (**Figure 1B**).

Since TagBFP can only be expressed in these cells via free Gal4p-VP64 released after SynNotch activation, we assessed functionality by assaying TagBFP expression as a function of Spike concentration (**Figure 1C, Supplementary Figure 1D**). We quantitatively characterized activation using mixture models to separate active cells from inactive cells in our cytometry histograms (**Figure 1D**, schematized in **Supplemental Figure 1C**, see Methods for detail). Cells expressing only the output transcriptional circuit but not SynNotch (No Notch) showed no change in expression in response to increasing Spike concentrations. To our surprise, ACE2-Notch was not appreciably activated even at large Spike concentrations, despite showing higher basal activation than the No Notch background. Remarkably however, LCB1-Notch showed robust activation, with 5-fold increase in mean TagBFP expression with as little as 0.3μg/ml Spike, a 20-50-fold increase for 1 and 3.16μg/ml Spike, and a slight decay in activation with a 20-fold increase in expression at 10μg/ml Spike. With its useful response characteristics, we dubbed LCB1-Notch *SARSNotch* and pursued its characterization further.

For sentinel cells to be useful therapeutics, it’s important that the sensing circuitry works in cells that are amenable to use for *in vivo* therapies. Traditional SynNotch designs targeting toy antigens such as GFP as well as blood tumor antigens like CD19 have been thoroughly tested in therapeutically relevant cells and in animal models^10,32,33^. However, none of these previous SynNotch variants have included a *de novo*-designed antigen sensor. To ensure SARSNotch functions in therapeutically relevant cells, we transduced primary human CD4+ T lymphocytes (T Cells) with the output circuit and SARSNotch and assessed SARSNotch function in the purified spike assay. T Cells expressing SARSNotch showed a similar dose range of activation, albeit with higher basal activation in comparison to Jurkats (**Figure 1E** and **1F, Supplementary Figure 1E**). These data establish SARSNotch’s potential utility in cell-based therapeutics. Further, it confirms that *de novo* designed protein binders may be a promising modular class of antigen sensors for SynNotch.

### SARSNotch can detect Spike protein with high sensitivity when surface expressed in a model of infected cells

We next sought to test SARSNotch activation in the presence of infected cells. Although the majority of Spike in infected cells is packaged into virions, Spike escapes along the biosynthetic pathway to be cellsurface displayed^34–36^. To model these infected cells with Spike on the cell surface, we generated a line of K562 cells expressing the perfusion stabilized Spike ectodomain^37^ tethered to the PDGFR transmembrane domain (Spike-K562s, **Figure 2A, Supplementary Figure 2A** and **2B**). We then co-cultured these cells at varying densities with SARSNotch-expressing Jurkat sentinels (**Supplementary Figure 2C**) and again assessed SARSNotch activation via TagBFP expression.

**Figure 2:**
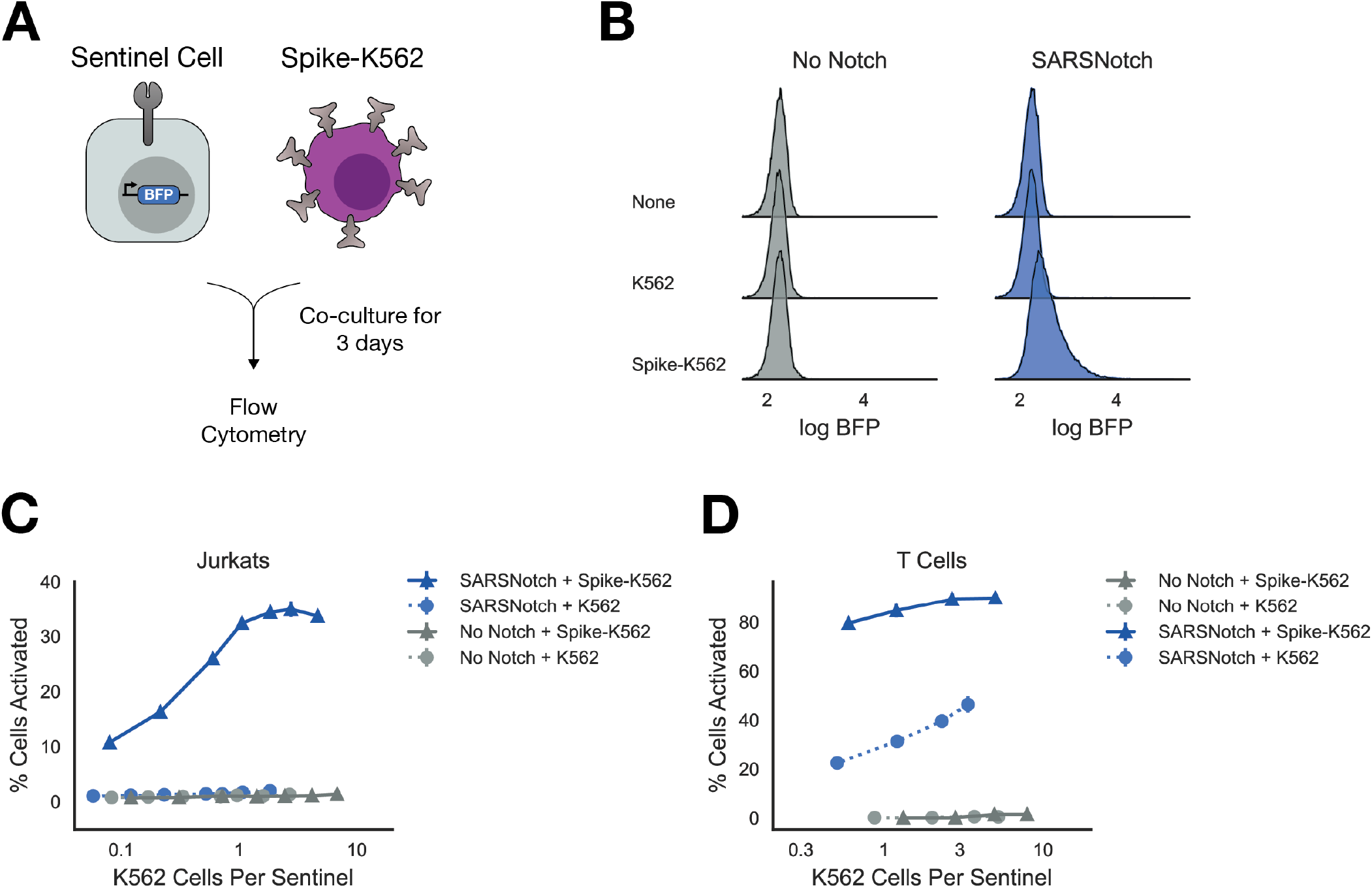
SARSNotch can detect Spike protein with high sensitivity when surface expressed in a model of infected cells. A) Schematic of experimental setup. K562 sender cells expressing the Spike ectodomain on their surface, modeling infected cells that express Spike on the cell surface, were co-cultured with sentinel cells expressing SARSNotch and BFP downstream of SARSNotch activation. B) Density estimates for Jurkat sentinel cell activation after co-culture with no, untransformed, or Spike-expressing K562 cells at equal density for 72 hours. C) Percent of activated Jurkat sentinel cells as a function of the ratio of K562 cells per sentinel cell after 72 hours of co-culture. D) Percent of activated sentinel T-cells as a function of the ratio of K562 cells per sentinel after 72 hours of co-culture. Data points and error bars represent mean and standard deviation of 3 biological replicates.

SARSNotch sentinel cells detected the presence of Spike-K562s (**Figure 2B**) at relative concentrations as low as 0.1 Spike-K562s per sentinel and reached saturation around 1 Spike-K562 per sentinel (**Figure 2C** and **Supplementary Figure 2E**). To assess the kinetics of the cell-sensing response, we plated Spike-K562s at approximately 1 Spike-K562 cell per sentinel and then assayed for activation at 24, 48, and 72 hours. We saw 20% of cells activated after 24 hours, increasing to 60% activated at 72 hours, suggesting a relatively rapid and sustained response of sentinels to the presence of Spike expressing cells (**Supplementary Figure 2D**).

We next tested SARSNotch T cells in the same cell-cell activation assay. T Cell SARSNotch sentinels showed a strong response to Spike-K562 cells even at low densities of Spike-K562 cells (**Figure 2D**). All T Cell sentinels, even those expressing No Notch, showed some activation in response to co-culture with K562s (**Figure 2D, Supplementary Figure 2F**), but the specific activation seen in SARSNotch-expressing T Cell sentinels in response to Spike-K562s was at least 2-fold greater than the non-specific activation for all K562 plating densities. These data confirm the ability of T Cell sentinels to sense cell-expressed Spike protein.

### SARSNotch does not increase susceptibility to viral infection

We next wanted to confirm that SARSNotch sentinel cells are not themselves a target for viral infection. Sentinel susceptibility to infection could be a dramatic weakness for an *in vivo* therapeutic as it could affect therapeutic cell survival and therefore treatment half-life, or in worst case scenarios it could enable sentinels to serve as viral reservoirs instead of therapeutics.

To test infection susceptibility, we generated lentiviruses encoding luciferase that were pseudotyped with either Spike, vesicular stomatitis virus glycoprotein (VSVG), or no glycoprotein (**Figure 3A**). We then incubated dilutions of the pseudotyped virus with SARSNotch sentinels or as a control, with a HEK293T cell line stably expressing the SARS-CoV-2 coreceptors ACE2 and TMPRSS2 that are susceptible to SARS-CoV-2 infection (293T-Susceptible). After 72 hours, we assayed cells for luciferase activity in both SARSNotch sentinels and 293T-Susceptible cells to evaluate for susceptibility to infection.

**Figure 3:**
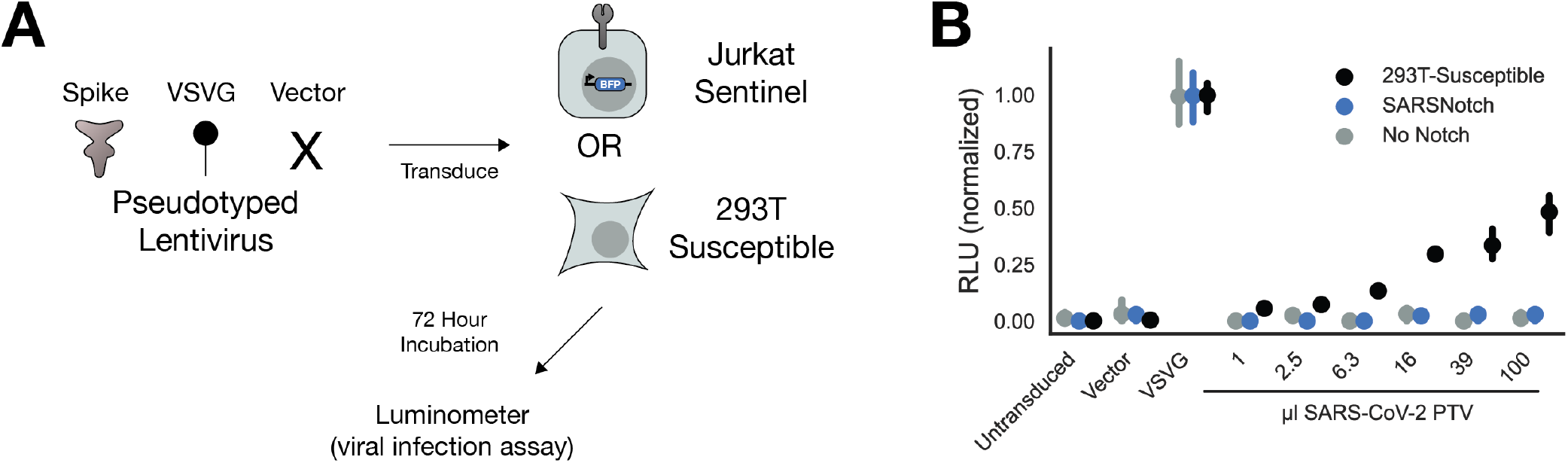
SARSNotch does not increase susceptibility to viral infection. A) Experimental schematic, showing that 3 different Luciferase-encoding pseudotyped virus variants were applied to either Jurkat sentinel cells or ACE2 and TMPRSS2-expressing HEK293T cells (293T-Susceptible), which were then assessed for SARSNotch activation and luciferase expression. B) Luciferase activity, normalized within each cell line to the maximum fluorescence from the VSVG infection, as a function of which pseudovirus was applied. Data points and error bars represent mean and 95% confidence interval of 3 biological replicates.

All cell types showed strong luminescence following infection with the positive control VSVG pseudovirus after 72 hours (**Figure 3B**, raw data in **Supplementary Figure 3A**), and even as early as 24 hours (**Supplementary Figure 3B** and **3C**). As expected, 293T-Susceptible cells were also infected in a dose dependent manner by the SARS-CoV-2 Spike pseudotyped virus. However, neither Jurkats alone nor our SARSNotch-expressing sentinels were susceptible to the SARS-CoV-2 Spike pseudotyped virus, showing no luminescence. This result demonstrates that our sentinel cells would not become viral reservoirs *in vivo*.

Although we designed SARSNotch sentinel cells mostly with the goal of detecting, and in future work acting on, infected cells displaying viral proteins, we also sought to determine whether viral particles (virions) themselves can be detected. Given the previously demonstrated inability of SynNotch to detect soluble ligands^28^, we hypothesized that SARSNotch would not be able to detect pseudotyped viral particles. Indeed, after incubating SARSNotch sentinels with pseudotyped virus as described above, we did not detect an increase in TagBFP expression for any dose of the pseudotyped virus at any time point assayed after infection (**Supplementary Figure 4A** and **B**). Like SARSNotch, ACE2-Notch was also unable to detect virions (**Supplementary Figure 4C**).

### SARSNotch shows dramatic activation in adherent cells

In addition to their potential uses as cell therapies, we envision that sentinel cells can enable an array of technologies to combat future threats. The ability to produce customized genetic responses to binding events positions sentinel cells as a potentially important reagent to generate better binders against SARS-CoV-2 Spike sequence variants or to rapidly generate optimized binders for other viruses using directed evolution systems. As a proof of principle for this use case, we investigated the functionality of SARSNotch in BHK-21 cells, a cell type relevant for the rapid mammalian directed evolution system VEGAS^38^.

To explore this possibility, we established a line of BHK-21 sentinels expressing SARSNotch, as well as the transcriptional output circuit (**Supplementary Figure 5**). We assayed SARSNotch activation in both purified protein and cell-cell assays for Spike detection as before. Unlike in Jurkat sentinels but similar to T Cells, we saw leakiness in SARSNotch activation in the absence of stimulus (**Figure 4A)**. However, despite this increased basal leakiness, SARSNotch BHK-21 sentinels showed dramatic activation in the presence of Spike, without the slight decrease in activation seen at high doses in the SARSNotch Jurkat sentinel cells (**Figure 4B**). A similar pattern was observed in the cell-cell assay, with almost 100% activation seen when SARSNotch BHK-21 sentinels were co-cultured with Spike-K562s at equal density for 72 hours (**Figure 4C, 4D**). This behavior highlights the modularity of SARSNotch across different cellular contexts, and suggests that there are many as yet unidentified contexts where this system could be used.

**Figure 4:**
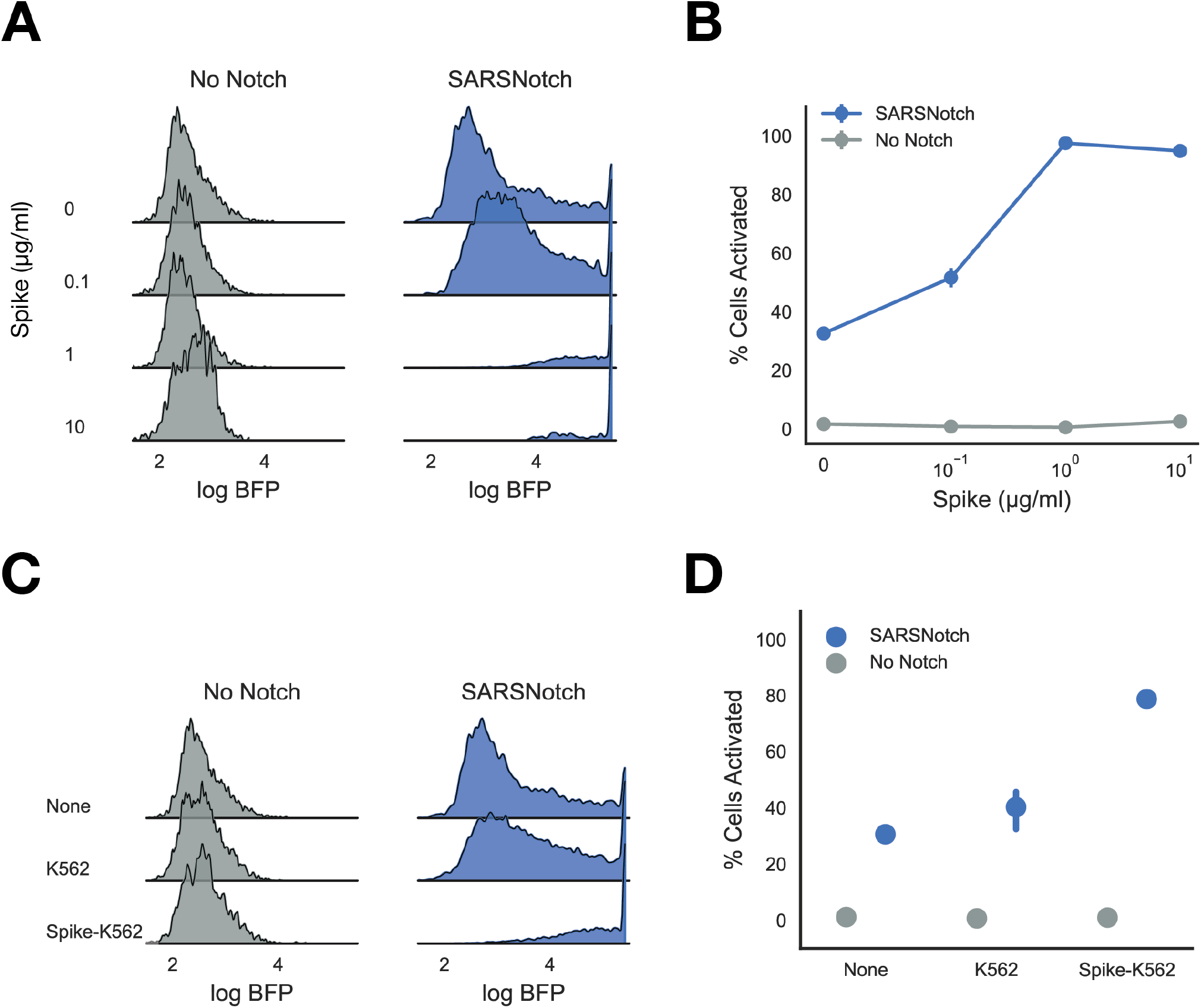
SARSNotch shows dramatic activation in adherent cell lines. A) Density estimates showing activation in No Notch and SARSNotch expressing BHK-21 cells as a function of purified Spike dose. B) Percent activated BHK-21 sentinel cells as a function of Spike dose. Quantification of data in Panel A. C) Density estimates showing activation in No Notch and SARSNotch BHK-21 sentinels as a function of cell-cell stimulation after co-culture with K562s at equal density for 72 hours. D) Percent activated sentinel BHK-21 cells for different sender cells (shown in panel C) used to activate them. Data points and error bars represent mean and standard deviation (B) or mean and 95% confidence interval (D) of 3 biological replicates.

## Discussion

In this work, we present SARSNotch, a SynNotch receptor coupled to a *de novo* designed SARS-CoV-2 Spike binder, which enables sentinel cells to detect Spike protein on its own or when surface displayed by opposing cells. Sensing is achieved within 24 hours of antigen presentation, and is sensitive to 0.3μg/ml purified Spike. Activation of SARSNotch also robustly occurs when Spike-expressing cells are present at as little as 1/10th the density of sentinel cells. Crucially, SARSNotch expression is not sufficient to increase the susceptibility of sentinel cells to SARS-CoV-2 viral infection. This sensing approach is portable between different cell types, including primary human T lymphocytes and adherent cells. This system therefore has the potential to deliver therapeutic payloads in a live cell-based therapy. Additionally, it can be used in therapeutic development schemes *in vitro*.

Selecting an antigen sensor to target a novel pathogen with SynNotch presents a unique challenge. All the SynNotch receptors described so far in the literature rely on either single chain variable fragment antibodies (scFvs) or nanobodies to recognize their antigen. Although both now exist against SARS-CoV-2 Spike^22,39^, no such reagents were available when we embarked on this project. We first speculated that the endogenous binder of Spike, ACE2, might be usable as an antigen sensor. The ACE2-Spike interaction is estimated to have a K_D_ ranging between 1.2 and 22nM^24,26,40,41^, whereas the scFv used to target the original SynNotch to CD19 has a K_D_ of 0.3nM^42^ and the scFv successfully used to adapt SynNotch to target HBV has a K_D_ of ~0.35nM^29^. However, we suspect this affinity difference is not the main reason ACE2-Notch didn’t yield a detectable signal output in the presence of Spike, given that an scFv against the human protein Her2 with K_D_ of 210nM has been recently described to enable SynNotch activation^43^. Perhaps more likely, the 600 amino acid-long ACE2 extracellular domain is between 2 and 3 times the size of the scFvs and nanobodies that have been used as antigen sensors previously, suggesting potential limits on the size of antigen sensors that SynNotch can accommodate. However, neither of these explanations are sufficient to explain ACE2-Notch’s deficit given the properties of LCB1, the *de novo*-designed binder that was successfully incorporated into SARSNotch. LCB1, and its partner lead candidate LCB3, are both 55 amino acids long with binding affinities below 1nM, yet only LCB1 lead to strong SynNotch activation downstream of Spike binding, whereas LCB3 produced only modest activation even at high doses (**Supplementary Figure 6**). This suggests that in addition to size, specific binding geometry may be required for SynNotch activation. Given these findings, we would still recommend a varied approach in developing future SynNotches with novel binders, exploring a wide range of proteins as antigen sensors. We also envision studies that use libraries of nanobodies, different proteins truncations, and synthetic antigen binders with different sizes, structures, and affinities to dissect the bio-mechanical properties of SynNotch.

An ideal cell-based therapy against a viral infection would 1) lack any propensity to be infected by the virus itself, lest it become a reservoir for the virus to replicate in, 2) detect cells that have succumbed to viral infection, and 3) detect virions before infection has occurred. SARSNotch meets both of the first two requirements, but falls short in the third. We suspect that sentinel cells expressing SARSNotch are not susceptible to SARS-CoV-2 because expression of both a viral receptor and a specific cell surface protease are important for SARS-CoV-2 infection^44^. It is also possible that LCB1 binds Spike in an orientation that does not enable cell entry, although ACE2-Notch also does not confer infection susceptibility to otherwise unsusceptible cells (**Supplementary Figure 3D)**. The inability of SARSNotch to detect virions is puzzling, especially given the previous adaptation of SynNotch to sensing HBV^29^. SynNotch, like Notch itself, is suspected to require force applied to the extracellular domain in order to activate, but insufficient force generated by the virions themselves seems unlikely given that both SARS-CoV-2 and the lentiviral pseudotyped virus used in this study produce virions ~100nm in diameter, twice the size of HBV particles ^45–48^. We suspect that in the case of the anti-HBV SynNotch, activation was due to presentation of virions in trans, either as they were bound to a hydrophobic tissue culture plate or by cells in the process of getting infected. If this is true, our SARSNotch sentinel cells may be more effective *in vivo* than our current data might suggest.

Regulated production of a genetically encoded therapeutic payload opens many possibilities for combating viral threats. Upon detection of infected cells, sentinel cells could produce and release virus-neutralizing proteins such as monoclonal antibodies^49^, multivalent nanobodies^22^, or even soluble versions of the *de novo*-designed binders used by the cells to detect the threat in the first place^23^. Such programs enable local delivery of otherwise systemically applied therapies. However, the power of sentinels lies in their ability to detect multiple signals and integrate them to produce even more specific responses than can be achieved through standard therapies. For example, longitudinal studies of COVID-19 patients have demonstrated immune misfiring present in severe cases that is absent in moderate cases^50^. Advanced sentinel cells could combine sensors like SARSNotch with synthetic sensors of endogenous immune signals^51–53^, using cellularly-implemented boolean logic^33^ to modulate immune function in an effort to mitigate disease severity.

Although the idea of using sentinel cells as cell-based therapeutics is enticing, significant work is necessary to develop them further for this end. Even if successful, cell-based therapies are presently costly and slow to generate. However, this may change in the future, as ongoing work generating allogeneic cell therapies shows significant promise^54^. We hope that this work, along with the prior art with HBV^29^ and HIV^55^, may lead to successful antiviral cell-based therapies in order to combat future threats, much as work on mRNA vaccines starting in the 1990s is proving useful to us today. In the meantime, these systems could be adapted for other use cases. Our specific interest is positioning SARSNotch, with its genetic gate, upstream of directed evolution systems. SARSNotch could be adapted to a system like VEGAS^38^, where a highly mutagenic virus could produce SARSNotch mutants that could be selected for higher binding affinity to Spike or pseudotyped virions. Alternatively, novel therapeutic compounds could be screened in SARSNotch cell-cell assays, where loss of fluorescent output from sentinel cells could identify compounds blocking SARSNotch and Spike interactions. Beyond these ideas, we suggest that there might be other uses of this anti-SARS-CoV-2 system that we have not yet envisioned.

To this end, with this manuscript we extend a community invitation for others to tailor this system to their own ideas, or collaborate with us to push this system forward. The COVID-19 pandemic has demonstrated the value of rapid scientific communication and collaboration. In sharing this, we hope that our tools can contribute to such innovation. We also suggest that continuing to extend cell-based therapies beyond cancer, and potentially in the realm of virology, has tremendous potential. We encourage you, the reader, to help us push the boundaries of what is possible as we build a new set of tools to combat disease.

## Methods

### Cell Culture

Jurkat and K562 cell lines were cultured in RPMI-1640 media (Gibco #11875-093) supplemented with 10% fetal bovine serum (FBS) (GE Healthcare #SH30071.03IH25-40), and 1% Anti-Anti (Gibco #15240-096). Primary human CD4+ T cells were isolated from anonymous donor blood after apheresis by negative selection (STEMCELL Technologies #15062 and 15023). T cells were cryopreserved in RPMI-1640 (Corning #10-040-CV) with 20% human AB serum (Valley Biomedical, #HP1022) and 5% DMSO (Sigma-Aldrich #472301). After thawing, T cells were cultured in human T cell medium consisting of X-VIVO 15 (Lonza #04-418Q), 5% Human AB serum and 10 mM neutralized N-acetyl L-Cysteine (Sigma-Aldrich #A9165) supplemented with 30 units/mL IL-2 (NCI BRB Pre-clinical Repository). BHK-21 cells (ATCC CCL-10) were cultured in MEM α with nucleosides and GlutaMAX (Gibco #32571-036) supplemented with 5% FBS, 1% Anti-Anti, and 10% Tryptose Phosphate Broth (Gibco #18050-039). HEK293T cells expressing ACE2 and TMPRSS2, a generous gift of Hannah S Sperber and Dr. Satish Pillai, were cultured in DMEM High Glucose (Gibco #10569-010) supplemented with 10% FBS, 1% of Antibiotic-Antimycotic, 10μg/ml Blasticidin S HCl, and 2μg/ml Puromycin. All cells were maintained at 37°C with 5% CO2

### Source of Primary Human T cells

Blood was obtained from Blood Centers of the Pacific (San Francisco, CA) as approved by the University Institutional Review Board. Primary CD4+ T cells were isolated from anonymous donor blood after apheresis as described above.

### Construct Preparation

The output circuit, pHR_Gal4UASpyb-TATA_tBFP_pGK_mCitrine, was a generous gift from Dr. Kole Roybal. All other plasmids were constructed using the Mammalian Toolkit (MTK)^31^, a hierarchical DNA assembly method based on Golden-Gate (GG) cloning^56^. New binding domains were domesticated into the MTK by ordering primers or gBlocks from IDT that contained the desired coding sequence with all BsaI and BsmBI restriction sites replaced with synonymous mutations and overhangs for MTK Part 3as, then ligated into the MTK part entry vector using a BsmBI GG reaction. For LCB1 and LCB3, protein sequences were translated to human optimized coding sequences using Benchling. The ACE2 extracellular domain (AA 1-615) was selected based on the sufficiency of this region for Spike binding^26^. The Spike ectodomain used for presentation in sender cells was based on a previously described structure-stabilized Spike variant^37^. All binding domains were domesticated with the CD8a signal sequence, a Myc tag, and a 2xG4S linker at their N termini (MALPVTALLLPLALLLHAARP-EQKLISEEDL-GGGGSGGGGS). Transcriptional units as described in **Supplementary Figures 1A and 2A** were assembled into a lentiviral destination backbones. All plasmids were propagated in Stbl3 *E. coli* (QB3 MacroLab). Domestication was verified via sequencing and transcriptional unit assembly was verified via restriction digest. All plasmids used in this study are available on request, and key plasmids will be available on Addgene.

### Generation of Stable Cell Lines

Lenti-X 293T cells (Takara Bio #632180) were seeded at approximately 7e5 cells/well in a 6-well plate to yield ~80% confluency the following day. The following day cells were transfected with 1.5μg of transfer vector containing the desired expression cassette, and the lentiviral packaging plasmids pMD2.G (170ng) and pCMV-dR8.91 (1.33μg) using 10μl of Lipofectamine 2000 (Invitrogen #11668-027) according to manufacturer protocols. Media was changed after 24 hours, and at 48 hours the viral supernatant was filtered through a 0.45μm PVDF filter and added to Jurkat or K562 cells seeded at approximately 1e5 cells/well in a 12-well plate. The plates with virus were then spun at 800xg for 45 minutes at 32C. After 24 hours, the viral media was then exchanged for fresh media. Polyclonal cell populations were selected via fluorescence-activated cell sorting (FACS) as described below. The HEK293T cell line expressing ACE2 and TMPRSS2 was generated as previously described^57^.

### Primary T cell Lentiviral Transduction

For each construct tested, 1e6 Primary T cells were thawed and activated the next day using CD3/CD28 Dynabeads (Life Technologies #11131D) at a 1:3 cell:bead ratio. Lentivirus was generated as above, but before cell application lentivirus was concentrated using the Lenti-X Concentrator kit (Takara Bio #631232) according to manufacturer’s instructions. Concentrated virus was applied directly to T cells (2 wells of virus per each SynNotch construct, 1 well of virus for output circuit). 24 hours after the addition of the viral supernatant to T cell culture, viral media was removed and fresh media was added. At day 5 post T cell stimulation, Dynabeads were removed and the T cells were sorted for construct expression via FACS. Sorted cells were maintained at 5e5 cells/ml until used for experiments.

### Antibodies

Surface expressed proteins were assayed for using Alexa Fluor 647 Anti-Myc tag antibody (Cell Signaling Technologies #2233S). Jurkat activation was assessed using Alexa Fluor 647 anti-CD25 (Biologend #302618) and BUV395 anti-CD69 (BD Biosciences #564364). All antibodies were diluted 1:100 in DPBS (UCSF Cell Culture Facility) for staining.

### FACS

Cell lines were bulk sorted for high expression using the UCSF Laboratory for Cell Analysis Core Facility FACSAriaII (BD Biosciences). Cells were assessed for mCitrine (488nm excitation, 530/30nm emission, 505lp collection dichroic), mCherry (561nm ex, 610/20nm em, 600lp cd), and Alexa647 (633nm ex, 670/30nm em) fluorescence. Fluorescence-negative controls were used to set detector power so that negative cells appeared to have mean fluorescence ~100 counts, and then transduced cells were sorted for cells with expression outside of the negative control expression level. All data were collected using FACSDiva (BD Biosciences)

### Flow Cytometry

Flow cytometry was performed using a LSR-Fortessa (BD Biosciences). Prior to the flow cytometry, cells were seeded at densities described below in a 96 well plates, using flat-bottom plates (Falcon #353072) for experiments involving BHK-21 cells and U bottom plates for all other experiments (Falcon #877217) and incubated for 24-72 hours as specified by the experiment. Plates were then spun down to settle all cells. For adherent cells, cells were incubated with 100μl Versene (Gibco #15040-066) for 3 minutes, then triturated and transferred to a U bottom plate. Plates were then decanted, and rinsed with DPBS (UCSF Cell Culture Facility). Cells were then resuspended and stained for 45 minutes with appropriate antibodies. After incubation, plates were spun down, rinsed with DPBS, and resuspended in flow buffer (DPBS with 10% FBS). The plates were then run on the flow cytometer using a four laser configuration (355nm, 405nm, 488nm, 561nm, 640nm), collecting fluorescence for BUV395 (355nm ex, 379/28 em), TagBFP (405nm ex, 450/50 em), mCitrine (488nm em, 530/30 em, 505lp cd), mCherry (561nm ex, 610/20 em, 600lp cd), and Alexa Fluor 647 (640nm ex, 670/14 em). At least 10,000 events were recorded for all single cell line assays, and 30,000 events for experiments involving two cell lines. All data were collected using FACSDiva (BD Biosciences)

### Spike Protein Expression and Purification

ExpiCHO-S cells (6E6cells/mL) were transfected with 1ug/mL of Spike protein DNA^22^ using the ExpiCHO™ Expression System Kit (Gibco #A29133) and shaken at 37°C and 8% CO2 in a 50mL culture. 18 hours posttransfection, cells were supplemented with ExpiCHO™ Feed and ExpiFectamine™ CHO Enhancer and shaken at 32°C and 8% CO2 for up to 14 days. Cells were spun at 1000xG for 10 minutes and the supernatant was kept to collect secreted Spike protein. Supernatant was transferred to 50 mL centrifuge tubes and supplemented with one Pierce protease inhibitor tablet (Thermo). The sample was then mixed with 2 mL washed Ni-Excel resin (Cytiva), and placed on a rocker for 1 hour for binding. After binding, the sample was washed with 25 bed volumes of Wash buffer (10 mM Tris/HCl, pH 8.0, 200 mM NaCl, 10 mM imidazole), and eluted with 7 bed volumes of elution buffer (10 mM Tris pH 8.0, 200 mM NaCl, 500 mM imidazole) into separate 2 mL centrifuge tubes. The elution was concentrated, filtered, and subjected to size-exclusion chromatography (Superose 6 10/300 Increase, Cytiva) using equilibration buffer (10 mM HEPES pH 8.0, 200 mM NaCl). All steps were performed at room temperature except for the chromatography which was done at 4C. The protein concentration was estimated based on the protein absorbance at 280nm with a spectrophotometer (Nanodrop One, Thermo), flash frozen, and stored in −80 °C.

### Purified Spike Activation Assay

An aliquot of purified Spike trimer protein in sterile filtered equilibration buffer (10 mM HEPES pH 8.0, 200 mM NaCl) was thawed on ice. Dilutions between 0.01 and 10μg/ml were prepared in equilibration buffer and 30μl of each was added per well of a 96 well plate. The plate was then spun down for 4 minutes at 400xg before being covered with foil, sealed, and put in a 4C refrigerator overnight. The following day, the protein-coated plate was then decanted and coated wells were rinsed with DPBS. At least 16 hours after plating 1.25e4 cells were plated in each well of the protein-bound plate prior to spinning at 400xg for 4 minutes. Cells were incubated with Spike for 72 hours and then assayed for activation via flow cytometry as described above.

### Cell-Cell Activation Assay

Sentinel cells expressing SynNotch variants were seeded at 1.25e4 cells/well in a 96-well plate. At the same time, K562 cells expressing indicated surface-targeted antigens were seeded at varying ratios to the sentinels, as indicated in **Supplementary Figure 2C**. At 24, 48, and 72 hours after plating, cells were assessed for activation. Due to cell line differences in proliferation, initial seeding densities were not consistent for each condition at the end points. At the end points, actual K562 density per sentinel cells was calculated by fitting a 2 component Gaussian mixture model to the distribution of mCitrine fluorescence to recover the actual number of sentinels (high mCitrine from the output circuit) vs. K562s (low mCitrine) per condition. All data are plotted as this final empirically-derived density.

### Pseudotyped Virus Production and Titer Quantification

SARS-CoV-2 spike typed pseudoviruses were generated as previously described with modifications^58^. Briefly, 293T cells were transfected with plasmid DNA (340 ng of Spike vector, 1μg CMV-Gag-Pol (pCMV-dΔR8.91), 125 ng pAdvantage (Promega), 1 μg Luciferase reporter (per 6-well plate)) for 48 h. For positive control, Spike vector was replaced with pMD2.G and for negative control this vector was omitted. Supernatant containing pseudovirus particles was collected, filtered (0.45μm), and stored in aliquots at −80°C. Pseudotyped viruses were quantified with a p24 assay (Takara Bio #632200) per manufacturer’s instructions.

### Pseudotyped Virus Assay

The day before infection, 1.25e4 293T cells expressing ACE2 and TMPRSS2, or Jurkats were seeded in a 96-well flat bottomed plate. On the day of infection, a dilution series of SARS-CoV-2 pseudotyped virus was added to the plated cells. Virus with equal titer to the highest dose used of the SARS-CoV-2 pseudotyped virus, as determined by p24 assay above, from the VSVG and vector pseudotyped controls was added to control wells. After incubation for 24 or 72 hours, cells were split in half for use in activation assay as described above, or for luminescence assay as described below.

### Luminescence Assay

Cells for were transferred to opaque-bottomed white plates (Grenier-Bio #655073) and then assessed for luminescence (Promega #E1501) per manufacturer’s protocol. Briefly, cells were lysed for 5 minutes and then incubated with luciferase substrate. Luciferase output was measured using a TECAN Infinite m1000 PRO with an integration time of 1s and either no attenuation (24 hours) or OD2 attenuation (72 hours).

### Data Presentation, Analysis, and Availability

All experiments were performed in at least biological triplicate. All data points are the mean of 3 replicates, with error presented as either standard deviation or ±95% confidence intervals where indicated. For flow cytometry experiments, means were calculated for all cells within a single replicate, and the presented means and error are calculated between replicates. Events collected from flow cytometry were filtered to remove small events and then density gated on FSC and SSC to capture singlet populations. Where presented, density estimates represent all events for all 3 replicates in a flow cytometry experiment, and are calculated via seaborn’s kdeplot function with bandwidth adjustment of 0.2. To calculate activation, conditions were first assessed for bimodality via exploratory data analysis. For experiments where no bimodality was observed, activation is expressed as mean fluorescence intensity of BFP. For experiments where bimodality was apparent, a 2 component Gaussian mixture model was fit to data pooled from all conditions to identify the means of the on and off components. Then percent activation was calculated by scoring the component memberships of all cells in a given condition. A similar process was used to calculate cell densities, using mCitrine fluorescence. All data analysis was conducted using custom Python scripts, available on github (https://github.com/weinberz/sarsnotch). Analysis was conducted in Jupyter^59^ and relied on numpy^60^, matplotlib, seaborn, pandas, SciPy^61^, scikit-learn^62^ and fcsparser.

## Author Contributions

ZYW and HE-S conceived of the project. ZYW, CEH, MK, and HE-S designed experiments. ZYW, CEH, and MK designed and created DNA constructs used in the study. ZYW, CEH, and MK conducted cell culture based experiments. LC and DB conceived of and designed the *de novo* binders. QCRG Structural Biology Consortium, represented by its protein purification team consisting of DD, EH, JL, SP, KS, AMS, FZ, generated purified Spike protein. MK generated pseudotyped virus and provided protocols for its use as overseen by RK and MO. SE and JG sorted cell lines used in this study.

## Acknowledgements

The authors would like to acknowledge Dr. Raymond Liu, Iowis Zhu, and Dr. Kole Roybal for discussion and advice on implementing SARSNotch. We thank Dr. Aashish Manglik for thoughtful discussion and manuscript review. We thank Dr. Colin Zamecnick and Dr. John Pak for assistance in pioneering the purified Spike assay. We thank Dr. Andrew H. Ng for suggestions on assay design for the purified Spike assay and overall advice. We thank Dr. Wendell A. Lim for the generous donation of the materials required for T Cell experiments. We thank Jesslyn Park and Dr. Nina Riehs for manuscript review. ZYW thanks Lindsey Osimiri for essential discussion, Stephanie E. Crilly for discussion and manuscript review, and Alex Stoitsiadis *et al*. and Hayley Williams *et al*. for essential assistance.

## References

1. Weber, E. W., Maus, M. V. & Mackall, C. L. The Emerging Landscape of Immune Cell Therapies. Cell 181, 46–62 (2020).

2. Hyrenius-Wittsten, A. & Roybal, K. T. Paving New Roads for CARs. Trends Cancer 5, 583–592 (2019).

3. Lim, W. A. & June, C. H. The Principles of Engineering Immune Cells to Treat Cancer. Cell 168, 724–740 (2017).

4. Viaud, S. et al. Switchable control over in vivo CAR T expansion, B cell depletion, and induction of memory. Proc. Natl. Acad. Sci. 115, E10898–E10906 (2018).

5. Liu, D., Zhao, J. & Song, Y. Engineering switchable and programmable universal CARs for CAR T therapy. J. Hematol. Oncol.J Hematol Oncol 12, 69 (2019).

6. Weber, E. W. et al. Pharmacologic control of CAR-T cell function using dasatinib. Blood Adv. 3, 711–717 (2019).

7. Fedorov, V. D., Themeli, M. & Sadelain, M. PD-1– and CTLA-4–Based Inhibitory Chimeric Antigen Receptors (iCARs) Divert Off-Target Immunotherapy Responses. Sci. Transl. Med. 5, 215ra172 (2013).

8. Chakravarti, D. & Wong, W. W. Synthetic biology in cell-based cancer immunotherapy. Trends Biotechnol. 33, 449–461 (2015).

9. Israni, D. V. et al. Clinically-driven design of synthetic gene regulatory programs in human cells. bioRxiv 2021.02.22.432371 (2021) doi:10.1101/2021.02.22.432371.

10. Roybal, K. T. et al. Engineering T Cells with Customized Therapeutic Response Programs Using Synthetic Notch Receptors. Cell 167, 419–432.e16 (2016).

11. Glasgow, A. A. et al. Computational design of a modular protein sense-response system. Science 366, 1024–1028 (2019).

12. Pan, X. et al. Expanding the space of protein geometries by computational design of de novo fold families. Science 369, 1132–1136 (2020).

13. Chen, Z. et al. De novo design of protein logic gates. Science 368, 78–84 (2020).

14. Lajoie, M. J. et al. Designed protein logic to target cells with precise combinations of surface antigens. Science 369, 1637–1643 (2020).

15. Langan, R. A. et al. De novo design of bioactive protein switches. Nature 572, 1–23 (2019).

16. Ng, A. H. et al. Modular and tunable biological feedback control using a de novo protein switch. Nature 572, 265–269 (2019).

17. Polack, F. P. et al. Safety and Efficacy of the BNT162b2 mRNA Covid-19 Vaccine. N. Engl. J. Med. 383, 2603–2615 (2020).

18. Baden, L. R. et al. Efficacy and Safety of the mRNA-1273 SARS-CoV-2 Vaccine. N. Engl. J. Med. 384, 403–416 (2021).

19. Sahin, U., Karikó, K. & Türeci, Ö. mRNA-based therapeutics — developing a new class of drugs. Nat. Rev. Drug Discov. 13, 759–780 (2014).

20. Crawford, E. D. et al. Rapid deployment of SARS-CoV-2 testing: The CLIAHUB. PLOS Pathog. 16, e1008966 (2020).

21. Abbott, T. R. et al. Development of CRISPR as an Antiviral Strategy to Combat SARS-CoV-2 and Influenza. Cell 181, 865–876.e12 (2020).

22. Schoof, M. et al. An ultrapotent synthetic nanobody neutralizes SARS-CoV-2 by stabilizing inactive Spike. Science 370, 1473–1479 (2020).

23. Cao, L. et al. De novo design of picomolar SARS-CoV-2 miniprotein inhibitors. Science 370, 426–431 (2020).

24. Walls, A. C. et al. Structure, Function, and Antigenicity of the SARS-CoV-2 Spike Glycoprotein. Cell S0092867420302622 (2020) doi:10.1016/j.cell.2020.02.058.

25. Yan, R. et al. Structural basis for the recognition of SARS-CoV-2 by full-length human ACE2. Science 367, 1444–1448 (2020).

26. Wrapp, D. et al. Cryo-EM structure of the 2019-nCoV spike in the prefusion conformation. Science 367, 1260–1263 (2020).

27. Pallesen, J. et al. Immunogenicity and structures of a rationally designed prefusion MERS-CoV spike antigen. Proc. Natl. Acad. Sci. 114, E7348–E7357 (2017).

28. Morsut, L. et al. Engineering Customized Cell Sensing and Response Behaviors Using Synthetic Notch Receptors. Cell 164, 780–791 (2016).

29. Matsunaga, S. et al. Engineering Cellular Biosensors with Customizable Antiviral Responses Targeting Hepatitis B Virus. iScience 23, 100867 (2020).

30. Ede, C., Chen, X., Lin, M.-Y. & Chen, Y. Y. Quantitative Analyses of Core Promoters Enable Precise Engineering of Regulated Gene Expression in Mammalian Cells. ACS Synth. Biol. 5, 395–404 (2016).

31. Fonseca, J. P. et al. A Toolkit for Rapid Modular Construction of Biological Circuits in Mammalian Cells. ACS Synth. Biol. 8, 2593–2606 (2019).

32. Srivastava, S. et al. Logic-Gated ROR1 Chimeric Antigen Receptor Expression Rescues T Cell-Mediated Toxicity to Normal Tissues and Enables Selective Tumor Targeting. Cancer Cell 35, 489–503.e8 (2019).

33. Williams, J. Z. et al. Precise T cell recognition programs designed by transcriptionally linking multiple receptors. Science 370, 1099–1104 (2020).

34. Duan, L. et al. The SARS-CoV-2 Spike Glycoprotein Biosynthesis, Structure, Function, and Antigenicity: Implications for the Design of Spike-Based Vaccine Immunogens. Front. Immunol. 11, (2020).

35. Hoffmann, M., Kleine-Weber, H. & Pöhlmann, S. A Multibasic Cleavage Site in the Spike Protein of SARS-CoV-2 Is Essential for Infection of Human Lung Cells. Mol. Cell 78, 779–784.e5 (2020).

36. Sadasivan, J., Singh, M. & Sarma, J. D. Cytoplasmic tail of coronavirus spike protein has intracellular targeting signals. J. Biosci. 42, 231–244 (2017).

37. Hsieh, C.-L. et al. Structure-based design of prefusion-stabilized SARS-CoV-2 spikes. Science 369, 1501–1505 (2020).

38. English, J. G. et al. VEGAS as a Platform for Facile Directed Evolution in Mammalian Cells. Cell 178, 748–761.e17 (2019).

39. Noy-Porat, T. et al. A panel of human neutralizing mAbs targeting SARS-CoV-2 spike at multiple epitopes. Nat. Commun. 11, 4303 (2020).

40. Lan, J. et al. Structure of the SARS-CoV-2 spike receptor-binding domain bound to the ACE2 receptor. Nature 581, 215–220 (2020).

41. Chan, K. K. et al. Engineering human ACE2 to optimize binding to the spike protein of SARS coronavirus 2. Science 369, 1261–1265 (2020).

42. Ghorashian, S. et al. Enhanced CAR T cell expansion and prolonged persistence in pediatric patients with ALL treated with a low-affinity CD19 CAR. Nat. Med. 25, 1408–1414 (2019).

43. Hernandez-Lopez, R. A. et al. T cell circuits that sense antigen density with an ultrasensitive threshold. Science 371, 1166–1171 (2021).

44. Hoffmann, M. et al. SARS-CoV-2 Cell Entry Depends on ACE2 and TMPRSS2 and Is Blocked by a Clinically Proven Protease Inhibitor. Cell 181, 271–280.e8 (2020).

45. Lamontagne, R. J., Bagga, S. & Bouchard, M. J. Hepatitis B virus molecular biology and pathogenesis. Hepatoma Res. 2, 163–186 (2016).

46. Shih, C., Yang, C.-C., Choijilsuren, G., Chang, C.-H. & Liou, A.-T. Hepatitis B Virus. Trends Microbiol. 26, 386–387 (2018).

47. Bar-On, Y. M., Flamholz, A., Phillips, R. & Milo, R. SARS-CoV-2 (COVID-19) by the numbers. eLife 9, e57309 (2020).

48. Vogt, V. M. & Simon, M. N. Mass Determination of Rous Sarcoma Virus Virions by Scanning Transmission Electron Microscopy. J. Virol. 73, 7050–7055 (1999).

49. Wang, C. et al. A human monoclonal antibody blocking SARS-CoV-2 infection. Nat. Commun. 11, 2251 (2020).

50. Lucas, C. et al. Longitudinal analyses reveal immunological misfiring in severe COVID-19. Nature 584, 463–469 (2020).

51. Schwarz, K. A., Daringer, N. M., Dolberg, T. B. & Leonard, J. N. Rewiring human cellular input–output using modular extracellular sensors. Nat. Chem. Biol. 13, 202–209 (2017).

52. Scheller, L., Strittmatter, T., Fuchs, D., Bojar, D. & Fussenegger, M. Generalized extracellular molecule sensor platform for programming cellular behavior. Nat. Chem. Biol. 14, 723–729 (2018).

53. Barnea, G. et al. The genetic design of signaling cascades to record receptor activation. Proc. Natl. Acad. Sci. U. S. A. 105, 64–69 (2008).

54. Perez, C., Gruber, I. & Arber, C. Off-the-Shelf Allogeneic T Cell Therapies for Cancer: Opportunities and Challenges Using Naturally Occurring “Universal” Donor T Cells. Front. Immunol. 11, (2020).

55. Maldini, C. R., Ellis, G. I. & Riley, J. L. CAR T cells for infection, autoimmunity and allotransplantation. Nat. Rev. Immunol. 18, 605–616 (2018).

56. Engler, C., Gruetzner, R., Kandzia, R. & Marillonnet, S. Golden Gate Shuffling: A One-Pot DNA Shuffling Method Based on Type IIs Restriction Enzymes. PLOS ONE 4, e5553 (2009).

57. Khanna, K. et al. Binding of SARS-CoV-2 spike protein to ACE2 is disabled by thiol-based drugs; evidence from in vitro SARS-CoV-2 infection studies. bioRxiv 2020.12.08.415505(2020) doi:10.1101/2020.12.08.415505.

58. Crawford, K. H. D. et al. Protocol and Reagents for Pseudotyping Lentiviral Particles with SARS-CoV-2 Spike Protein for Neutralization Assays. Viruses 12, 513 (2020).

59. Kluyver, T. et al. Jupyter Notebooks - a publishing format for reproducible computational workflows. in Positioning and Power in Academic Publishing: Players, Agents and Agendas (eds. Loizides, F. & Scmidt, B.) 87–90 (IOS Press, 2016).

60. Harris, C. R. *t al.* Array programming with NumPy. Nature 585, 357–362 (2020).

61. Virtanen, P. et al. SciPy 1.0: fundamental algorithms for scientific computing in Python. Nat. Methods 17, 261–272 (2020).

62. Pedregosa, F. et al. Scikit-learn: Machine Learning in Python. J. Mach. Learn. Res. 12, 2825–2830 (2011).

