## Supplementary Figures for "Sentinel cells enable genetic detection of SARS-CoV-2 Spike protein"

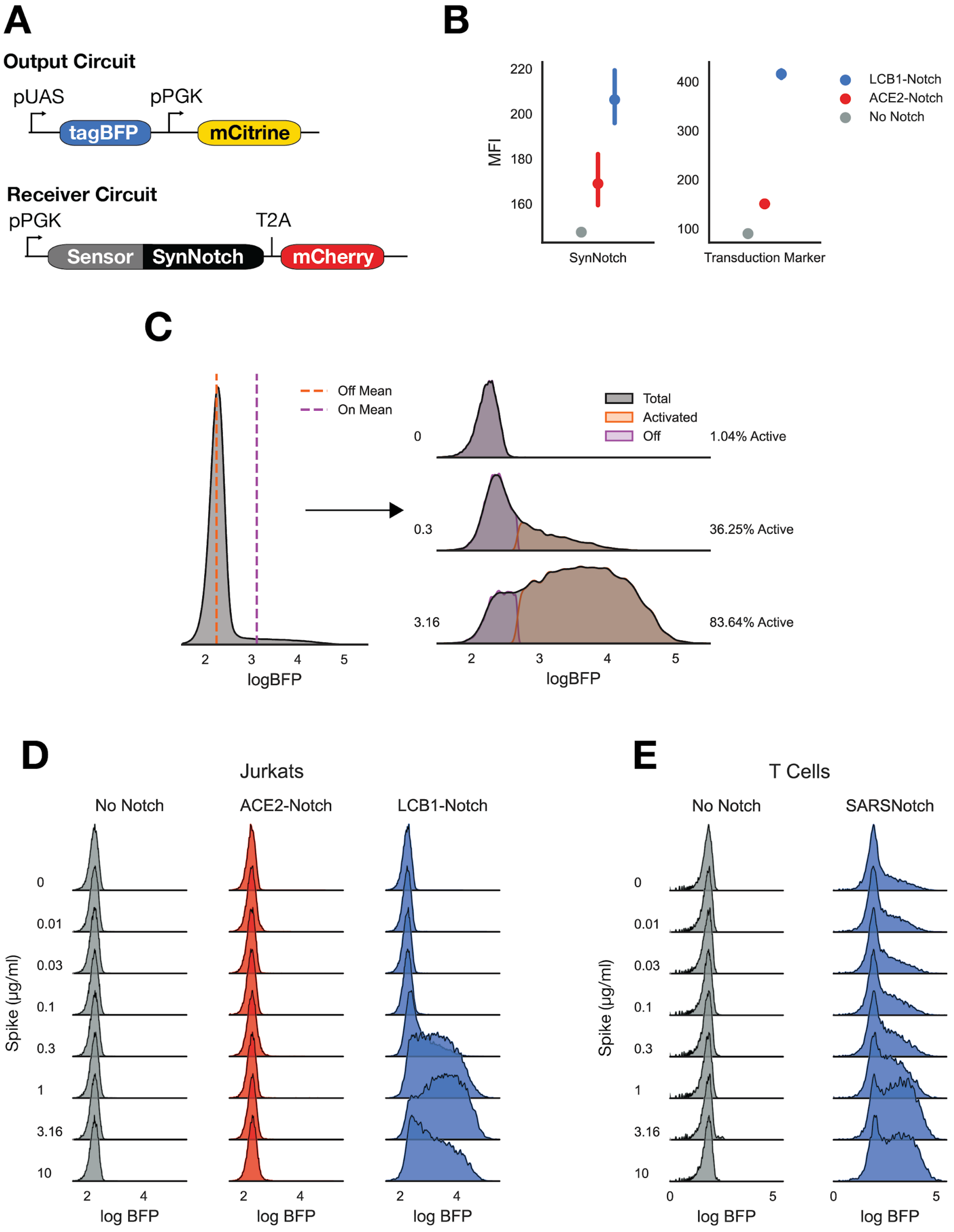

**Supplementary Figure 1: Sentinel cell line characterization, quantification, and alternative designs.** A) The two primary constructs used in all sentinel cells. The Output Circuit, with tagBFP expression downstream of the (5X)UAS ybTATA promoter, and a constitutively expressed mCitrine, was expressed in all sentinel cells used in this study. The Receiver Circuit, with swappable sensor domains for the different antigen sensors used in the study linked directly to the Notch Core and Gal4p-VP64 transcription factor, followed by a T2A element and mCherry expression serving as a transduction marker. B) Mean fluorescent intensity of SynNotch expression (left) as measured by anti-Myc antibody staining, and of the transduction marker mCherry (right) in the cell lines used in **Figure 1**. C) To quantify activation, a Guassian mixture model was fit to the logBFP distribution for all cells in an experiment pooled across conditions (left). To assess activation, the model was used to predict the fraction of events in each condition belonging to the ‘Active’ and ‘Off’ populations (right). D) Density estimates for Jurkat sentinel activation at all Spike doses tested. E) Density estimates for T Cell sentinel activation at all Spike doses tested. Data points and error bars represent mean and 95% confidence interval of 3 biological replicates.

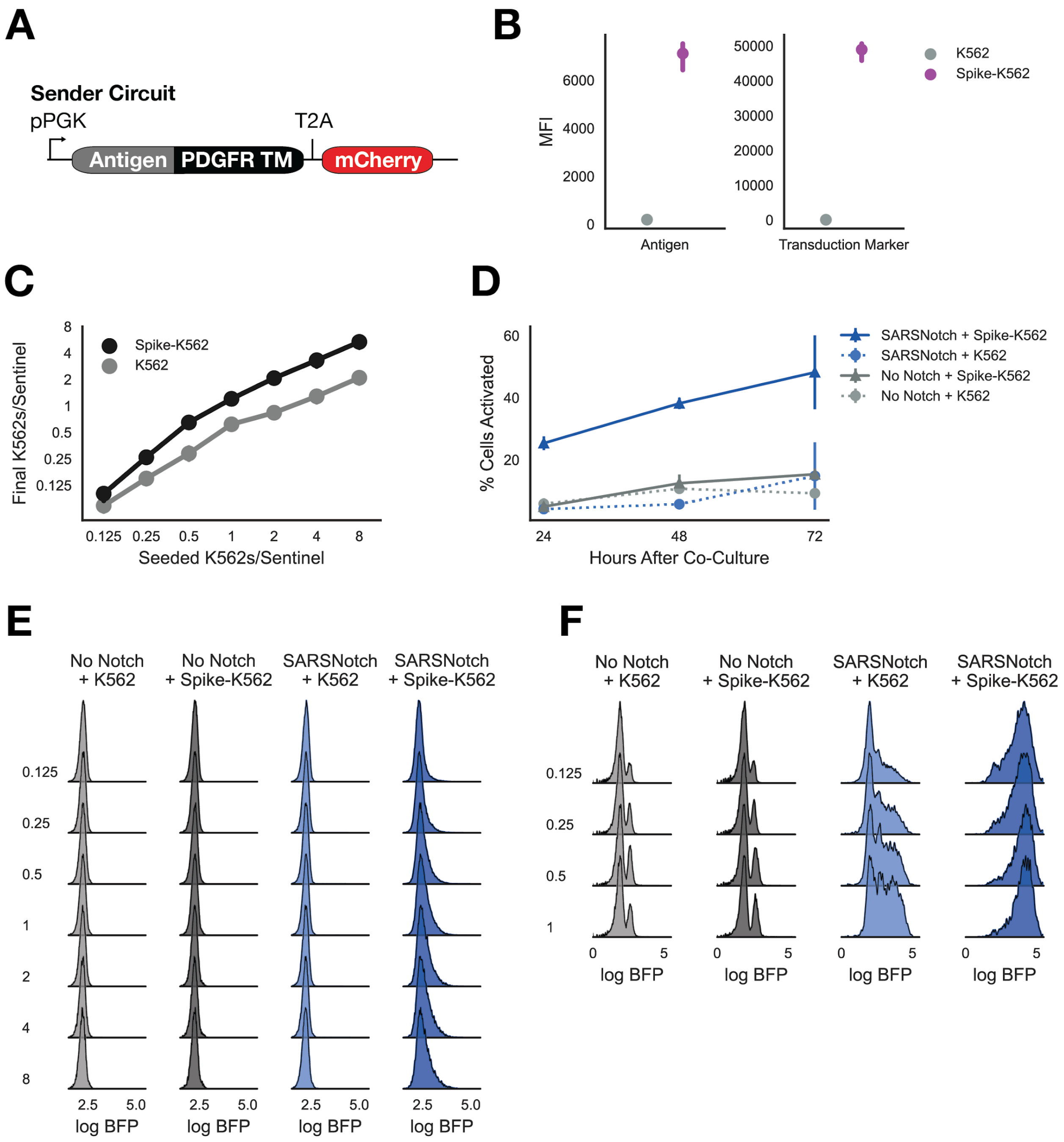


**Supplementary Figure 2**: **Characterization of sender cell lines**. A) The Sender Circuit places an extracellular domain directly N-terminal of the PDGFR transmembrane domain, followed by a T2A element and mCherry expressed as a transduction marker. B) Surface expression of the target antigen as assessed by anti-Myc staining (left) and transduction marker expression (right) for each of the sender K562 lines used in this study. C) For cell-cell experiments, cells were seeded to achieve desired K562 per sentinel densities (X axis). Due to differing rates of proliferation between cell lines, the final density at 72 hours differed slightly between cell lines (Y axis). D) Jurkat sentinel cell activation as a function of time after co-culture with K562s at equal density. E) Density estimates of activation in Jurkat sentinel cells for all K562 co-culture experiments. F) Density estimates of activation in T Cell sentinel cells for all K562 co-culture experiments. Data points and error bars represent mean and 95% confidence interval (B and C) or mean and standard deviation (D) of 3 biological replicates.

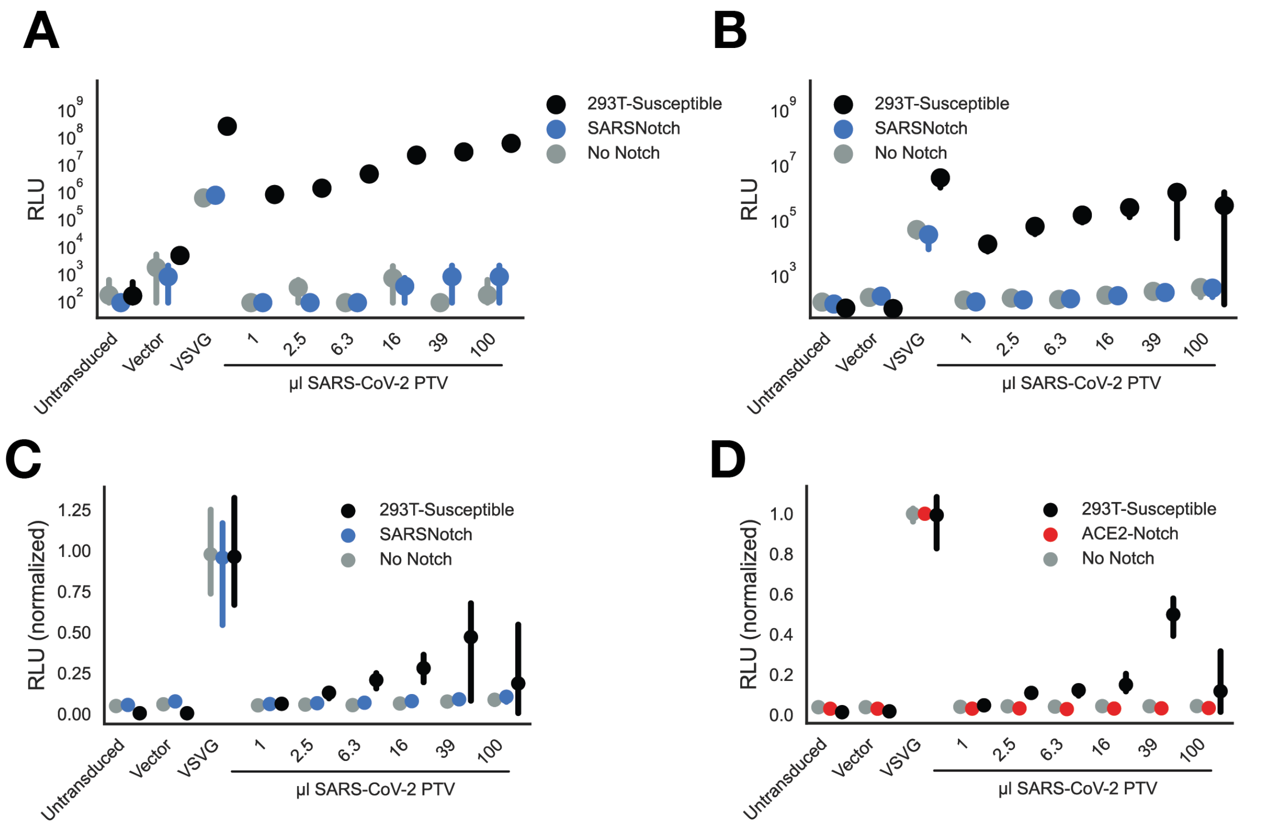


**Supplementary Figure 3: Supplementary luminescence data for pseudovirus assay**. Raw luminescence values as a function of pseudovirus application at 72 hours (A) and 24 hours (B). Normalized luminescence for SARSNotch Jurkat sentinels 24 hours after virus addition (C), or for ACE2-Notch Jurkat sentinels 72 hours after virus addition (D). Data points and error bars represent mean and 95% confidence interval of 3 biological replicates.

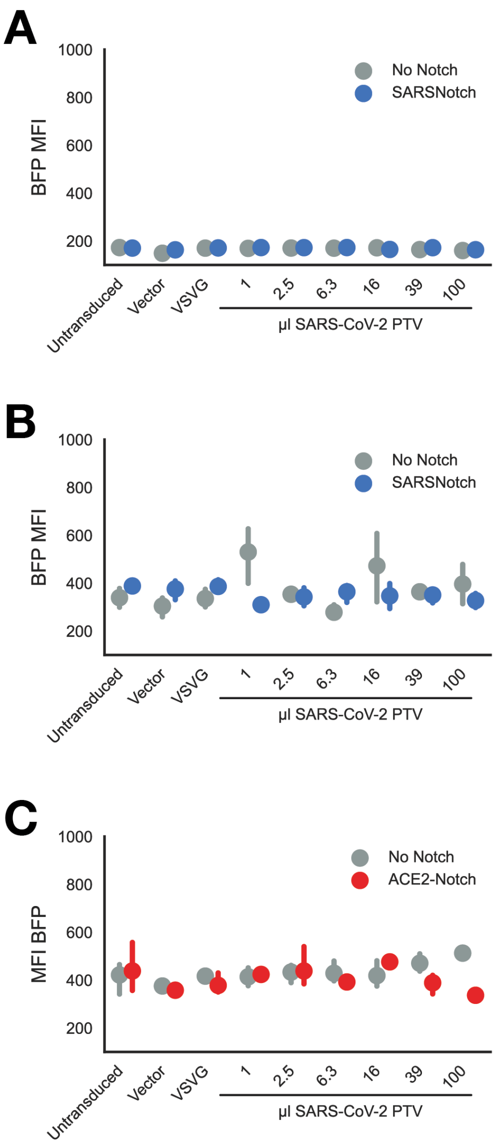


**Supplementary Figure 4: SARSNotch and ACE2-Notch activation in the presence of pseudovirus.** Activation assessed as mean fluorescence intensity (MFI) of BFP for each pseudotyped virus-treated Jurkat sentinel cell condition at 72 hours (A) or 24 hours (B) after infection, and for ACE2-Notch 72 hours after infection (C). Data points and error bars represent mean and 95% confidence interval of 3 biological replicates

**A**

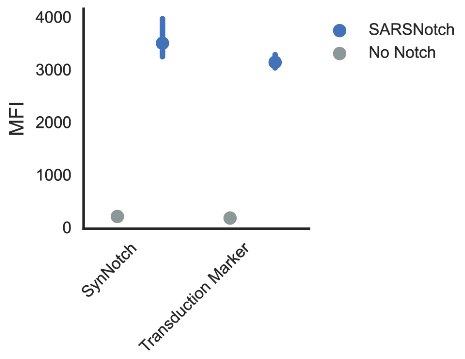


**Supplementary Figure 5: Characterization and activation quantification for BHK-21 sentinel cells.** A) SynNotch expression as assessed by anti-Myc staining, and transduction marker expression in BHK-21 No Notch parental line and SARSNotch-expressing line. Data points and error bars represent mean and 95% confidence interval of 3 biological replicates

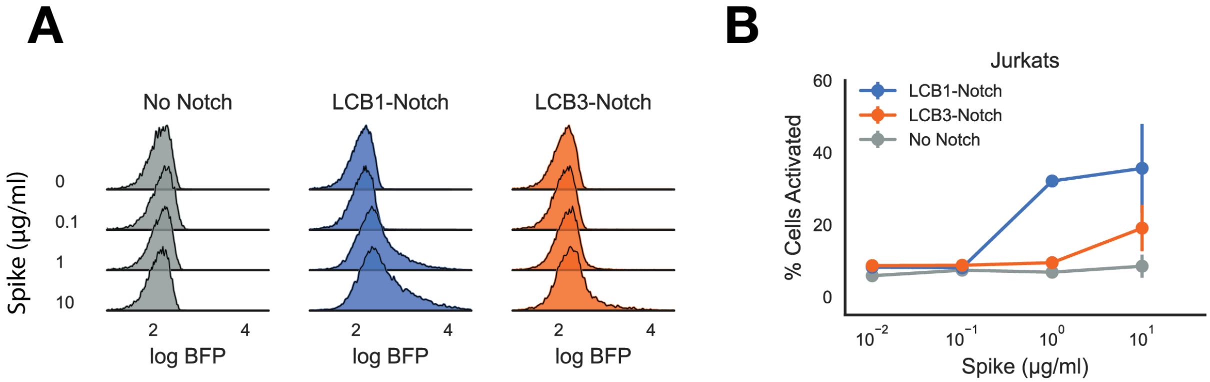


**Supplementary Figure 6: LCB1 is a better SynNotch antigen sensor than the next lead *de novo*-designed candidate LCB3.** A) Density estimates for activation of Jurkat sentinels expressing No Notch, LCB1-Notch (SARSNotch), or LCB3-Notch as a function of Spike protein dose. B) Quantified activation for sentinels in A as a function of Spike dose. Data points and error bars represent mean and standard deviation of 3 biological replicates.
